# Predicting longitudinal traits derived from high-throughput phenomics in contrasting environments using genomic Legendre polynomials and B-splines

**DOI:** 10.1101/632117

**Authors:** Mehdi Momen, Malachy T. Campbell, Harkamal Walia, Gota Morota

## Abstract

Recent advancements in phenomics coupled with increased output from sequencing technologies can create the platform needed to rapidly increase abiotic stress tolerance of crops, which increasingly face productivity challenges due to climate change. In particular, the high-throughput phenotyping (HTP) enables researchers to generate large-scale data with temporal resolution. Recently, a random regression model (RRM) was used to model a longitudinal rice projected shoot area (PSA) dataset in an optimal growth environment. However, the utility of RRM is still unknown for phenotypic trajectories obtained from stress environments. Here, we sought to apply RRM to forecast the rice PSA in control and water-limited conditions under various longitudinal cross-validation scenarios. To this end, genomic Legendre polynomials and B-spline basis functions were used to capture PSA trajectories. Prediction accuracy declined slightly for the water-limited plants compared to control plants. Overall, RRM delivered reasonable prediction performance and yielded better prediction than the baseline multi-trait model. The difference between the results obtained using Legendre polynomials and that using B-splines was small; however, the former yielded a higher prediction accuracy. Prediction accuracy for forecasting the last five time points was highest when the entire trajectory from earlier growth stages was used to train the basis functions. Our results suggested that it was possible to decrease phenotyping frequency by only phenotyping every other day in order to reduce costs while minimizing the loss of prediction accuracy. This is the first study showing that RRM could be used to model changes in growth over time under abiotic stress conditions.

## Background

Plant biology has become a large-scale, data-rich field with the development of high-throughput technologies for genomics and phenomics. This has increased the feasibility of data driven approaches to be applied to address the challenge of developing climate-resilient crops (Tester and Langridge, 2010). Crop responses to environmental changes are highly dynamic and have a strong temporal component. Such responses are also known as function-valued traits, for which means and covariances along the trajectory change continuously. Single time point measurements of phenotypes, however, only provide a snapshot, posing a series of challenges for research efforts aimed at understanding the ability of the plant to mount a tolerant response to an environmental constraint. Advancements in high-throughput phenotyping (HTP) technologies have enabled the automated collection of measurements at high temporal resolution to produce high density image data that can capture a plethora of morphological and physiological measurements (Furbank and Tester, 2011). In particular, image-based phenotyping has been deemed a game changer because conventional phenotyping is laborious and often involves destructive methods, precluding repeated sampling over time from the same individual (Ge et al., 2016). More importantly, these HTP systems offer greater potential to uncover the time-specific molecular events driven by important genes that have yet to be discovered in genome-wide association studies (GWAS) or to perform forecasting of future phenotypes in longitudinal genomic prediction. Thus, integrating these HTP data into quantitative genetics has the potential to increase the rate of genetic gain in crops. However, to take full advantage of such opportunities, novel statistical methods that can fully leverage time series HTP data need to be developed.

Recently, Campbell et al. (2018) used a random regression model (RRM) to perform genomic prediction for longitudinal HTP traits in controlled environments, such as greenhouses, using Legendre polynomials as the choice of a basis function to model dependencies across time. They also demonstrated that RRM could be used to achieve reasonable prediction accuracy in a cross-validation (CV) framework to forecast future phenotypes based on information from earlier growth stages. RRM also enables the calculation of (co)variances and genetic values at any time between the beginning and end of the trajectory, even including time points that lack phenotypic information. This study showed that RRM could effectively describe the temporal dynamics of genetic signals by accounting for mean and covariance structures that are subjected to change over time (Kirkpatrick et al., 1990). However, the utility of RRM for plants under an abiotic stress environment is not explored. This is a critical unknown as the crop productivity is greatly limited by environmental challenges such as drought and heat stress. In addition to the Legendre polynomials, spline functions can be used to describe the relationships between image-based phenomics and genomics data in longitudinal modeling (White et al., 1999). In particular, B-spline functions have been used to study a variety of traits, such as growth records, in animal breeding in terms of model goodness of fit using pedigree data (e.g., Meyer, 2005; Baldi et al., 2010); however, its application to HTP data in plants and its predictive ability from a CV perspective remains untested.

Here we present our results from the performance of RRM applied to HTP temporal shoot biomass data in response to control and water-limited conditions using Legendre polynomials and spline functions. We selected drought stress because water limitation significantly impacts shoot growth (PSA) and is the major limitation for agricultural productivity and global food security.

## Materials and Methods

### Plant materials and greenhouse conditions

Three hundred fifty-seven accessions (*n* = 357) of the rice (*O. Sativa*) diversity panel 1 (RDP1) were selected for this study (Zhao et al., 2011). Seeds were surface sterilized with Thiram fungicide and germinated on moist paper towels in plastic boxes for three days. For each accession, three uniformly germinated seedlings were selected and transplanted to pots (150mm diameter x 200 mm height) filled with 2.5 kg of UC Mix. Square containers were placed below each pot to allow water to collect. The plants were grown in saturated soil on greenhouse benches prior to phenotyping.

All lines were genotyped with 44,000 single nucleotide polymorphisms (SNPs) (Zhao et al., 2011). We used PLINK v1.9 software (Purcell et al., 2007) to remove SNPs with a call rate ≤ 0.95 and a minor allele frequency ≤ 0.05. Missing genotypes were imputed using Beagle software version 3.3.2 (Browning and Browning, 2007). Finally, 36,901 SNPs were retained for further analysis.

### Experimental design and drought treatment

All experiments were conducted at the Plant Accelerator, Australian Plant Phenomics Facility, at the University of Adelaide, SA, Australia. The panel was phenotyped for a digital metric representing shoot growth over 20 days of progressive drought using an image-based phenomics platform. The details of the experimental design are provided in Campbell et al. (2018). Briefly, each experiment consisted of 357 accessions from RDP1 and was repeated three times from February to April 2016. Two smart-houses were used for each experiment. In each smart-house, the accessions were distributed across 432 pots positioned across 24 lanes. The experiments followed a partially replicated paired design, where plants of the same accession were grown adjacent to one another. In each experiment, 54 accessions were randomly selected and replicated twice.

Seven days after transplant (DAT), plants were thinned to one seedling per pot. Two layers of blue mesh were placed on top of the pots to reduce evaporation. The plants were loaded on to the imaging system and were watered to 90% field capacity (FC) DAT. On the 13 DAT, each pot was watered to 90% and was imaged to obtain an initial phenotype before the onset of drought. One plant from each pair was randomly selected for drought treatment. Water was withheld from drought plants until 25% FC, and after which water was applied to maintain 25% FC. For the duration of the experiment, the control plants were maintained at 100% FC.

### Statistical analysis of phenotypic data

Visible images were processed, and digital features were extracted using the open-source Python library Image Harvest (Knecht et al., 2016). The sum of plant pixels from the two sides and one top view of red/green/blue (RGB) images was summed and used as a measure of shoot biomass. This digital phenotype is referred to as the projected shoot area (PSA) throughout this study. Several studies have reported a high correlation between PSA estimates and shoot biomass (Campbell et al., 2015; Golzarian et al., 2011; Knecht et al., 2016). Prior to downstream analyses, outlier plants at each time point were detected for each trait using the 1.5 interquartile range rule, and potential outliers were plotted along with their treatment counterparts and inspected visually. Plants that exhibited abnormal growth patterns were removed. In total, 221 plants were removed, leaving 2,586 plants for downstream analyses.

Raw phenotypic measurements were adjusted for downstream genetic analyses prior to fitting RRM. Best linear unbiased estimators (BLUE) were computed for each accession by fitting experimental effect with three levels and replication within experiment for some of the accessions. We postulated that observations at each time point follow the additive genetic model 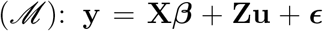, where **X** and **Z** are *n*’ × *f* and *n*’ × *n* orders of incident matrices linking observations (*n*’) to systematic effects (*f*) and number of accessions (*n*), respectively, **y** is an *n*′ × 1 vector of observations at each time point, ***β*** is a *f* × 1 vector of systematic effects, **u** is a *n* × 1 vector of BLUE for accessions, and e is an *n*^1^ × 1 vector of residuals with 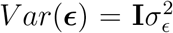, where **I** is an identity matrix. This was followed by fitting a RRM-based genomic prediction approach to predict phenotypes as described below.

### Random regression model

We conducted quantitative genetics modeling of image-derived phenotypes using a RRM to assess how well we could predict dynamic genetic signals. The RRM assumes that genetic effects and genetic variances are not constant and can vary continuously across the trajectory. This leads to better prediction of time-dependent complex traits by estimating heterogeneous single nucleotide polymorphism (SNP) effects across the trajectory. Specifically, we viewed the trajectory of digital image-processed longitudinal records as an infinite-dimensional characteristic that could be modeled by a smooth function (Meyer and Hill, 1997; Van der Werf et al., 1998). Changes in PSA over time were modeled through Legendre polynomials and B-splines of time at phenotyping. The general formula for the RRM was as follows:

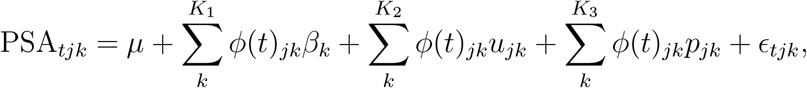

where *φ*(*t*)_*jk*_ is a time covariate coefficient defined by a basis function evaluated at time point *t* belonging to the *j*th accession; *β_k_* is a *k*th fixed random regression coefficient for the population’s mean growth trajectory; *u_jk_* is a *k*th random regression coefficient associated with the additive genetic effects of the *j*th accession; *K*_1_ is the number of random regression parameters for fixed effect time trajectories; *K*_2_ and *K*_3_ are the number of random regression parameters for random effects; and *p_jk_* is a *k*th permanent environmental random regression coefficient for the accession *j*. The starting values of index *k*, and *K*_1_, *K*_2_, and *K*_3_ are defined separately for Legendre polynomials and B-splines below.

In the matrix notation, the above equation can be rewritten as

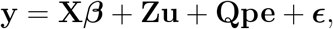

where ***β*** is a vector of solutions for fixed regressions; **u** is the additive genetic regression coefficients; **pe** is the permanent environmental random regression coefficients; ***ϵ*** is the residuals; and **Z** and **Q** are corresponding incident matrices. We assumed a multivariate-Gaussian distribution and the variance-covariance structure of

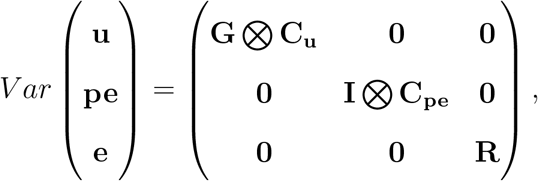

where 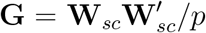 is the genomic relationship matrix of VanRaden (2008), where **W**_*sc*_ represents a centered and standardized marker matrix and *p* is the number of markers; **C**_*u*_ is the covariance function between the random regression coefficients for the additive genetic effect; ⊗ is the Kronecker product; **C**_*pe*_ is the covariance function between the random regression coefficients for the permanent environmental effects; and 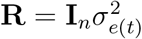 is a diagonal matrix of heterogeneous residuals varying across times, where 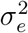 is the residual variance.

### Choice of basis function

The choice of the basis function to model the shape of the longitudinal measurements is critical. An ideal basis function has adequate potential to capture real patterns of changes in variance along a continuous scale (time) for a given trait (Meyer and Kirkpatrick, 2005). In this study, we used RRM with two basis functions, i.e., Legendre polynomials (Meyer, 1998) and B-splines (Meyer, 2005), to describe line-specific curves for the PSA trajectory over the day of imaging.

#### Legendre polynomials

Applying parametric shape functions for covariates of time is challenging because these covariates tend to generate high correlations among trajectories (Mrode, 2014). For this reason, fitting Legendre polynomials of time at recording as covariables is a common choice to model growth curves because these polynomials greatly reduce the correlations between estimated random regression coefficients and make no prior assumptions regarding the shape of the longitudinal curve. This function has been used widely in animal breeding for many years (e.g., Jamrozik and Schaeffer, 1997) and has recently been used in plant breeding as well (Sun et al., 2017; Campbell et al., 2018; Marchal et al., 2019). Suppose d is the order of fit or degree of the polynomial. Legendre polynomials evaluated at the standardized time points were computed as **Φ** = **MΛ**, where **M** is the *t_max_* by *d* +1 matrix containing the polynomials of the standardized time covariate 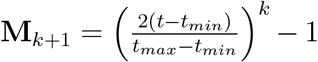 and Λ is the *d* +1 × *d* +1 matrix of Legendre polynomial coefficients (Kirkpatrick et al., 1990). Here, *t_min_* = 1 and *t_max_* = 20 because PSA was measured for 20 days. We chose the same orders of polynomials for fixed, additive, and permanent environmental coefficients as previously described Schaeffer (2016). We compared linear (*k* = 0, …, *K*_1_ = *K*_2_ = *K*_3_ = 1) and quadratic (*k* = 0, …, *K*_1_ = *K*_2_ = *K*_3_ = 2) Legendre polynomials in this study. Thus, the numbers of regression coefficients were *d* + 1 = 2 and *d* + 1 = 3 for linear and quadratic Legendre polynomials, respectively.

#### B-splines

Spline functions consist of individual segments of polynomials joined at specific points called knots. B-splines first require determination of the total number of knots K. Although a large number of knots will increase complexity, too few knots will decrease accuracy. This basis function is reported to offer several advantages, including better numerical properties compared with polynomials, especially when there are high genetic variances at the extremes of the trajectory period, negative correlations between the most distant time point measurements, and a small number of records, particularly at the last stage of the trajectory (Meyer, 2005; Misztal, 2006). Here, we used equidistant knots, and the B-spline function was computed from Cox-de Boor’s recursion formula (De Boor, 2001). Given a preconsidered knot sequence of time t, the covariables for B-splines of degree *d* = 0 were defined by assuming values of unity for all points in a given interval or zero otherwise. For the ith interval given by knots

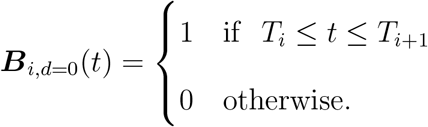

where *T* is the threshold in time interval. According to De Boor (2001), the matrix **Φ** of B-spline for higher-order polynomials can be defined by recursion

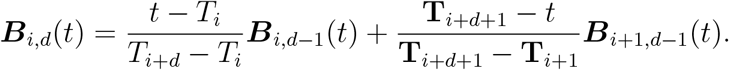

This indicates that a B-spline of degree *d* is simply a function of B-splines of degree *d* – 1. Note that the number of random regression coefficients depends on the number of knots and order of polynomials for B-splines. In general, the number of regression coefficients is given by *K* = *s* + *d* – 1 (Meyer, 2005). In this study, we fitted linear B-splines with *s* = 3 or *s* = 4 knots to divide the time points into equally spaced intervals. The same number of knots was considered for fixed trajectories, additive genetic, and permanent environmental coefficients. Thus, the numbers of regression coefficients were three (*k* =1, …, *K*_1_ = *K*_2_ = *K*_3_ = 3 + 1 – 1 = 3) and four (*k* = 1, …, *K*_1_ = *K*_2_ = *K*_3_ = 4 + 1 – 1 = 4) for *s* = 3 and *s* = 4 knots, respectively.

### Goodness of model fit

The goodness of fit of RRM was assessed by computing the Akaike’s information criterion (AIC) (Akaike, 1974) and the Schwarz—Bayesian information criterion (BIC) (Schwarz et al., 1978). The best model was selected based on the largest AIC and BIC values after multiplying by −1/2. We used Wombat software to fit RMM in this study (Meyer, 2007).

### Cross-validation scenarios

As graphically represented in Figure 1, three different CV scenarios were designed to train the RRM. In all scenarios, prediction accuracy was evaluated by computing Pearson correlations between predicted genetic values and PSA in the testing set. Each of the CV scenarios is described below.

**Figure 1:**
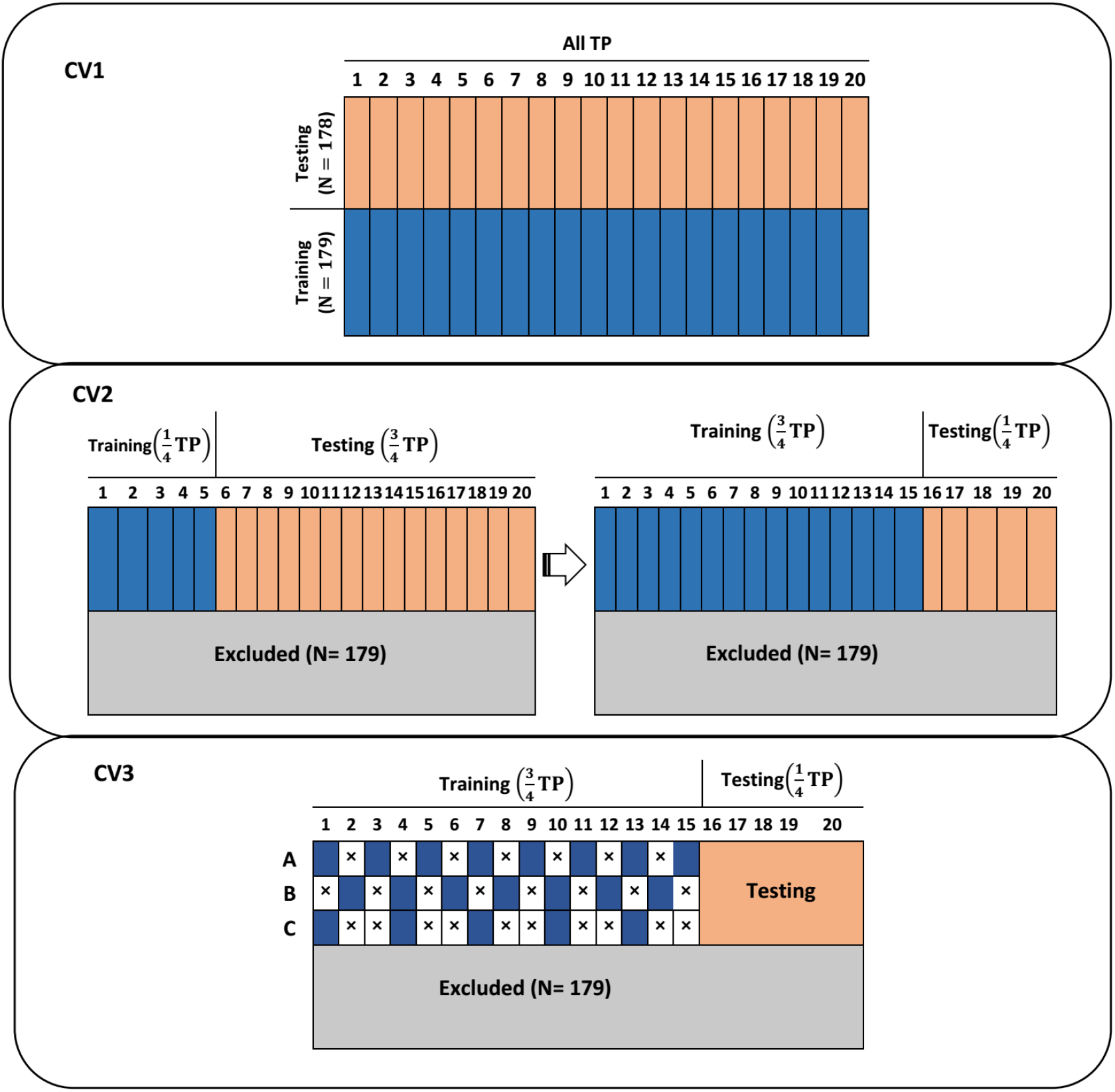
Pictorial representation of three cross-validation schemes used for predicting longitudinal projected shoot area (PSA) using a random regression model coupled with Legendre polynomials and B-splines. The data set consisted of 357 lines. CV1: 179 lines were used as the training set to predict PSA for the remaining 178 lines. Here, all 20 time points in the training set were used to predict PSA at each of 20 time points for a new set of lines. CV2: The data set was split into two longitudinal stages. The model was trained using the earlier growth stages to predict PSA at the second part of growth stages. We increased the number of time points used for training in a sequential manner. CV3: This was used to evaluate the impact of phenotyping frequency in the training data set on longitudinal prediction accuracy. Observations on odd days were used (CV3A), Observations on even days were used (CV3B), and keep one and delete two consecutive time points (CV3C). TP: time points.

#### CV1

In the first CV scenario (CV1), the whole data set was divided into two subsets, i.e., training and testing sets, each including 179 and 178 accessions, respectively. All 20 time points in the training set were fit to the RRM using Legendre polynomials and B-splines, and we predicted phenotypic values of 20 time points for lines in the testing set. Random assignment of individuals into the training and testing sets was repeated 10 times. The equation for CV1 was set up in the following manner. The time-specific genetic value of the ith individual in the training set was computed as 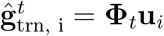, where 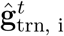 is the estimated genetic value of the individual *i* at time *t*; **Φ**_*t*_ is the *t*th row vector of the *t_max_* × *K*_1_ matrix **Φ**; and **u**_*i*_ is the *i*th column vector of the *K*_1_ × *n* matrix **u**. Then, a vector of predicted genetic values of individuals in the testing set at time *t* was obtained as 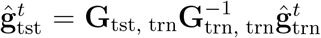, where **G**_tst, trn_ is the genomic relationship matrix between the testing and training set and 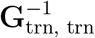 is the inverse of genomic relationship matrix between the training set. Because CV1 is not a forecasting task, a standard multi-trait model (MTM) was also fitted as a baseline model considering longitudinal traits as different traits (Henderson and Quaas, 1976). The BLUPF90 family of programs was used to fit MTM with 20 traits (Misztal et al., 2002).

#### CV2

The second CV scenario (CV2) was related to forecasting future phenotypes from longitudinal traits at early time points. Individuals in the training set were used to forecast their yet-to-be observed PSA values at later time points from information available at earlier time points. The first quarter of the time points {*t* = 1, 2, 3, 4, 5} was used as the training set, and the remaining time points {*t* = 6, 7,· · ·, 20} were predicted for each line in the training set. This was followed by sequentially increasing the number of time points used to train the model so that in the last run, three quarters of the time points {*t* = 1,2, … 15} were used in the training set to forecast phenotypes at the last quarter of time points {*t* = 16, 17, 18, 19, 20}. This CV scenario was designed to find a sufficient set of earlier time points to obtain reasonable longitudinal prediction accuracy and is known as walk forward validation. We set up the CV2 equation by first estimating the random regression coefficient matrix **u** using **Φ**_1:*t*_, which was computed from time point 1 to time point *t*. The prediction of future time points *t*’ (*t* +1 ≤ *t*’ ≤ *t*_max_) for an individual *i* was carried out by 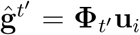, where **Φ**_*t*’_ is the *t*’th row vector of *t*_max_ – *t* by *K* +1 matrix **Φ**; and **u**_*i*_ is the *i*th column vector of the number of random regression coefficients by *n* matrix **u**.

#### CV3

The third CV scenario (CV3) was designed to evaluate whether it was possible to reduce the phenotyping frequency while still maintaining a high prediction accuracy for the last quarter of observations. We used the last case in CV2 such that time points {*t* = 1, 2, …, 15} were used for the training set to forecast the last quarter of observations {*t* = 16, 17, 18, 19, 20}. We then reduced the number of time points used in the training set as follows: A, observations on odd days {*t* =1,3, …, 15} were used; B, observations on even days {*t* = 2,4, …, 14} were used; C, keep one and delete two consecutive time points. In CV2 and CV3 scenarios, half of the individuals were randomly selected to fit the model, and the analysis was repeated 10 times. If the loss of prediction accuracy was minimal, it was possible to reduce the phenotyping cost. The equation for CV3 was set up in the same way as that for CV2.

### Data availability

Genotypic data regarding the rice accessions can be downloaded from the rice diversity panel website (http://www.ricediversity.org/). Phenotypic data used herein are available in Supplementary File S1.

## Results

### Assessing model fit

Figures 2A and 2B show the box plots of the original PSA and BLUE for the phenotypic trajectories over the 20 days of imaging for control and water-limited conditions. The PSA for control and water-limited plants diverged significantly after 10 days of initiation of the drought treatment, and the accession level difference become apparent at later growth stages under control conditions. Supplementary Figure 1 shows the linear or quadratic forms of Legendre polynomials and three and four knot-based B-spline curves over 20 days of imaging. For Legendre polynomials, intercept, linear, and quadratic coefficients are represented in black, red, and green, respectively. For B-spline, knot 1, knot 2, and knot 3 are represented in black, red, and green, respectively.

**Figure 2:**
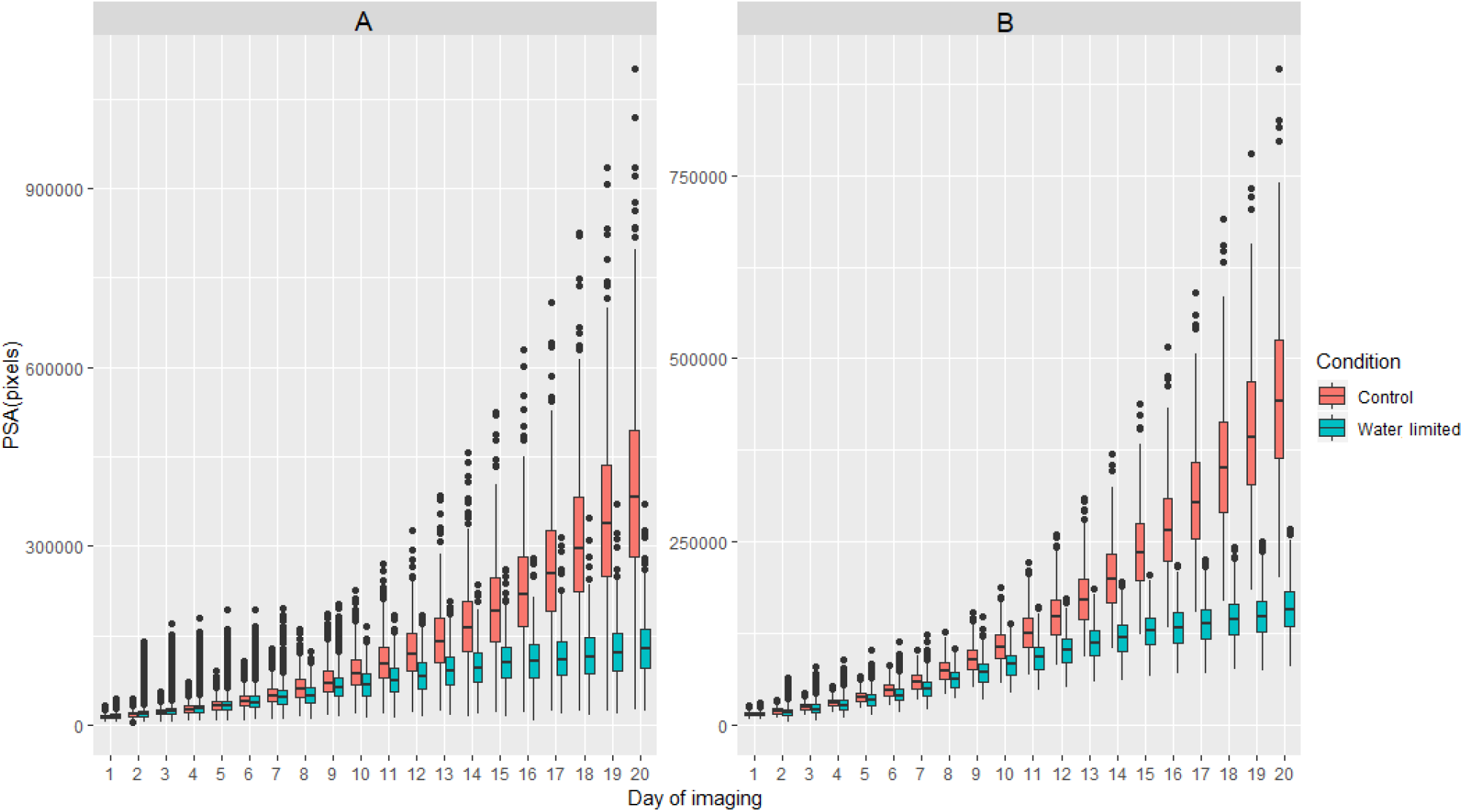
A: Box plots of projected shoot area (PSA) over the 20 days of imaging in two environments: controlled and water-limited conditions. B: Best linear unbiased estimators over the 20 days of imaging in two environments: controlled and water-limited conditions.

Table 1 summarizes the goodness of fits of RRM coupled with linear and quadratic Legendre polynomials and B-spline functions in control and water-limited conditions. For the Legendre polynomials, quadratic forms require more parameters to be estimated compared with linear forms. Similar to observation for B-splines, the presence of a greater number of knots suggested that there were more parameters to be estimated. Under control conditions, the best goodness of fit was obtained by linear Legendre polynomials, followed by linear B-splines with three knots, linear B-splines with four knots, and quadratic Legendre polynomials according to AIC scores. According to BIC scores, linear Legendre polynomials, followed by linear B-splines with three knots, quadratic Legendre polynomials, and linear B-splines with four knots. Under water-limited conditions, the best goodness of fit was given by linear Legendre polynomials, followed by linear B-splines with three knots, quadratic Legendre polynomials, and linear B-splines with four knots for both AIC and BIC scores. The number of parameters in the model varied from 26 to 40.

**Table 1:** Assessing goodness of fit for two random regression models (Legendre polynomials and B-splines) used to predict projected shoot area measured over 20 time points.

| Condition | CF | Log L | −0.5 AIC | −0.5 BIC | $p$ |
| --- | --- | --- | --- | --- | --- |
| CON | LEGL | −32414.493 | −32440.493 | −32529.839 | 26 |
|  | LEGQ | −32412.550 | −32444.550 | −32554.512 | 32 |
|  | BSPL3 | −32408.862 | −32440.862 | −32550.824 | 32 |
|  | BSPL4 | −32404.142 | −32444.142 | −32581.592 | 40 |
| WL | LEGL | −26011.867 | −26037.867 | −26127.213 | 26 |
|  | LEGQ | −26009.267 | −26041.267 | −26151.229 | 32 |
|  | BSPL3 | −26006.205 | −26038.205 | −26148.167 | 32 |
|  | BSPL4 | −26005.537 | −26045.537 | −26182.986 | 40 |

CON: control environment; WL: water-limited environment; CF: covariance function; LEGL: Legendre polynomial linear; LEGQ: Legendre polynomial quadratic; BSPL3: B-spline linear with three knots; BSPL4: B-spline linear with four knots; Log L: log likelihood; AIC: Akaike information criterion; BIC: Bayesian information criterion; and $p$ : number of parameters.

### Cross-validation

The results from CV1 are shown in Figure 3. This CV was designed to evaluate the accuracy of predicting testing set individuals using all time points. Under control conditions, MTM performed relatively better than RRM up to day 3. The prediction accuracy of RRM increased subsequently and after the 10th day of imaging, the best prediction was given by linear Legendre, followed by quadratic Legendre, linear B-spline with three knots, and linear B-spine with four knots. Overall, RRM performed better than MTM, and linear Legendre was the best prediction machine throughout the growth stages. Under water-limited conditions, prediction accuracy was lower compared with those of control conditions. All RRM delivered higher prediction than MTM except for the first two time points. Although Legendre polynomials performed better than B-splines until day 7, the difference between these approaches became negligible afterward.

**Figure 3:**
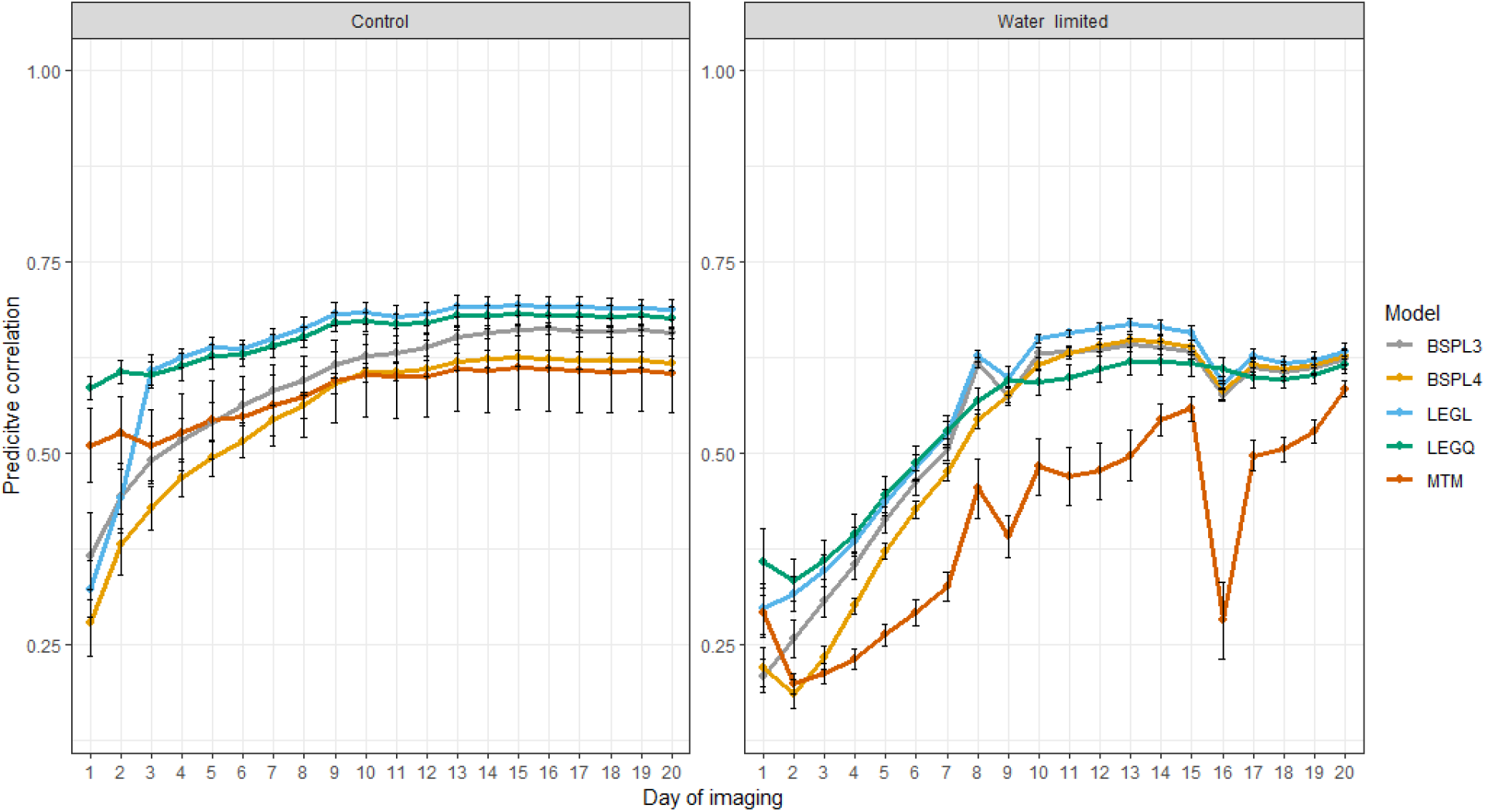
Prediction accuracy obtained from cross-validation 1 scenario. Total of 179 lines were used as the training set to predict PSA for the remaining 178 lines. Here, all 20 time points in the training set were used to predict PSA at each of 20 time points for a new set of lines. LEGL: linear Legendre polynomials; LEGQ: quadratic Legendre polynomials; BSPL3: linear B-splines with three knots; BSPL4: linear B-spline with four knots; MTM: multi-trait model.

Figures 4 and 5 show the accuracy of CV2 under control and water-limited conditions, respectively. This CV was designed to test how much information from the previous time points was required to achieve reasonable prediction accuracy at later growth stages. Under control conditions, we found that the best prediction for the last five time points was achieved when using all time point information up to the most recent (15/5 CV2 subscenario). This suggested that having more information from previous time points to train the model would result in higher prediction accuracy. Using the first five time points to train the model resulted in the worse prediction (5/15 CV2 subscenario). Thus, it is likely that the prediction accuracy in RRM declined because we attempted to estimate numerous parameters from only five time points. Legendre polynomials yielded better and more stable prediction than B-splines. We observed a similar trend under water-limited conditions; that is, using more previous time points to train the model resulted in higher prediction accuracy. However, the accuracy of prediction was unstable and decreased dramatically. There was no noticeable difference between the Legendre polynomials and B-splines in terms of performance.

**Figure 4:**
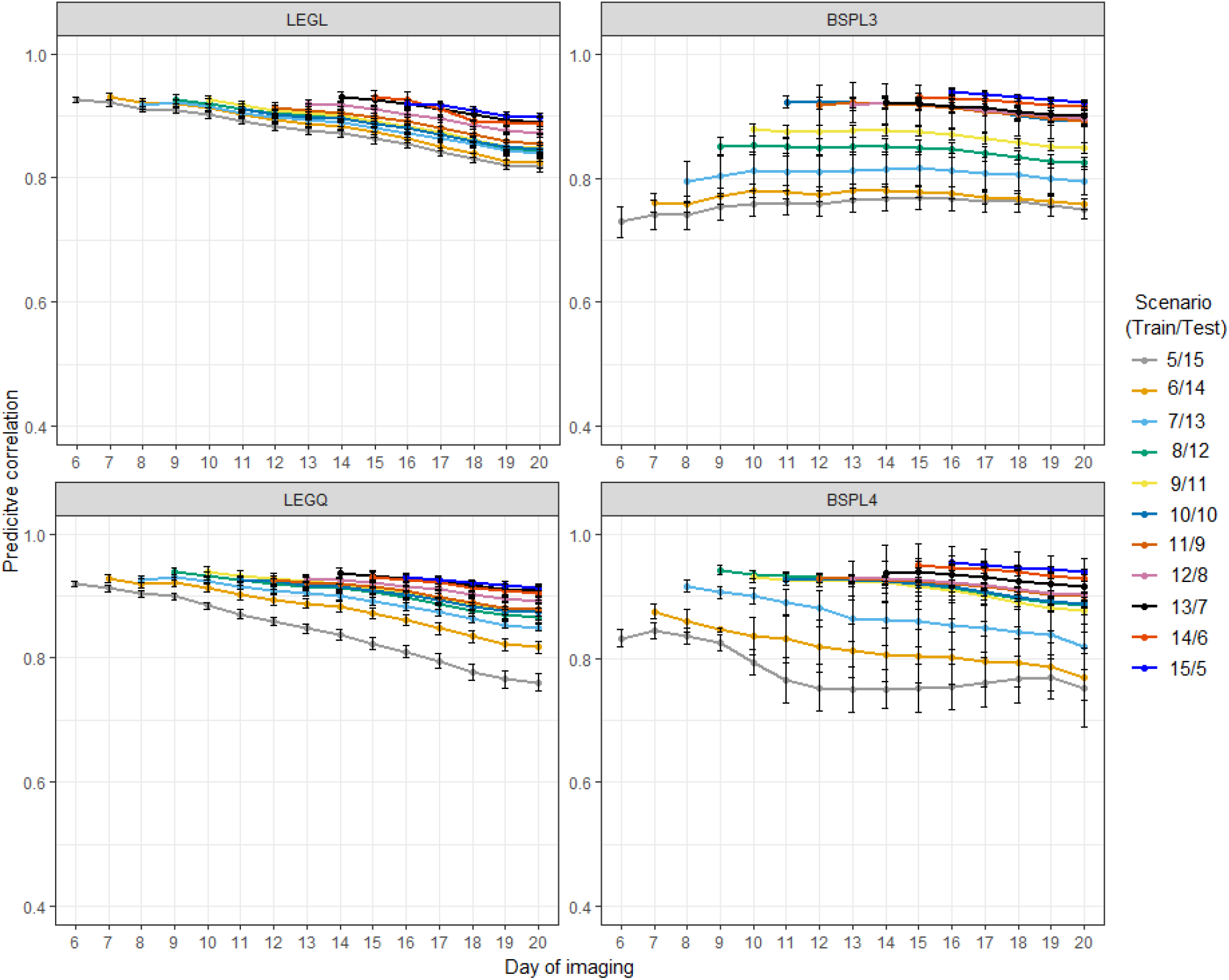
Prediction accuracy of cross-validation scenario 2 in control conditions. Each line depicts the different number of training and testing sets partitioning at the time point levels. LEGL: linear Legendre polynomials; LEGQ: quadratic Legendre polynomials; BSPL3: linear B-splines with three knots; BSPL4: linear B-spline with four knots.

**Figure 5:**
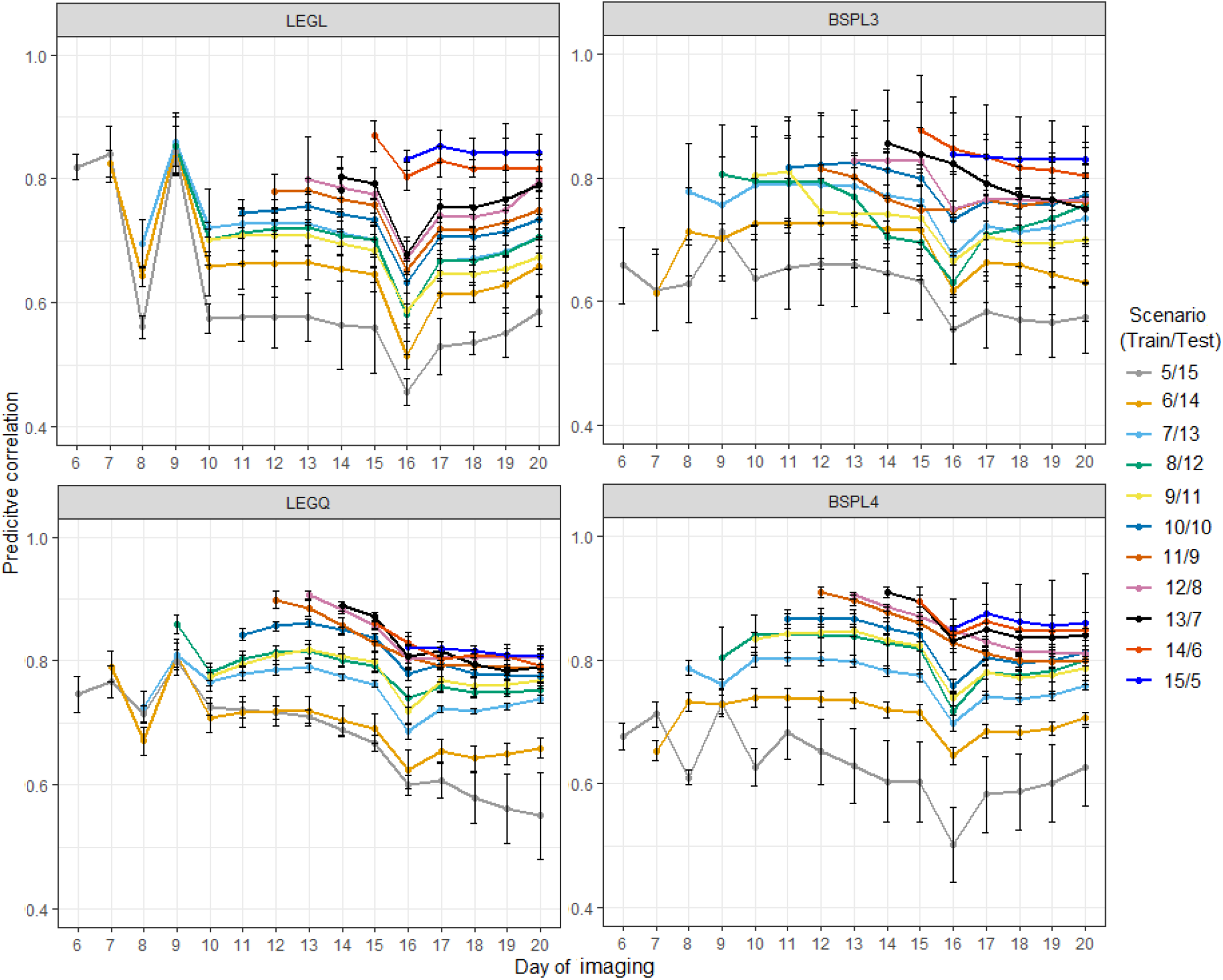
Prediction accuracy of cross-validation scenario 2 in water-limited conditions. Each line depicts the different number of training and testing sets partitioning at the time point levels. LEGL: linear Legendre polynomials; LEGQ: quadratic Legendre polynomials; BSPL3: linear B-splines with three knots; BSPL4: linear B-spline with four knots.

Figures 6 and 7 show the CV3 accuracy under control and water-limited conditions, respectively. We designed this CV to evaluate whether it was possible to reduce phenotyping frequency and phenotyping costs without sacrificing prediction accuracy. Under control conditions, the prediction accuracy of CV3A, CV3B, and CV3C all decreased relative to the benchmark scenario in CV2, where all of the first 15 time points were used for the training set to forecast the last five time points. Although removing two consecutive time points did not improve performance (CV3C), the prediction accuracy from phenotyping every other day was still relatively high (CV3A and CV3B). In general, the linear B-splines performed the best, and differences between scenarios were minimal. Under water-limited conditions, we observed the same trend, but the prediction accuracy was more unstable and decreased relative to control conditions. The quadratic Legendre polynomials and B-splines with four knots did not perform well, possibly due to overfitting.

**Figure 6:**
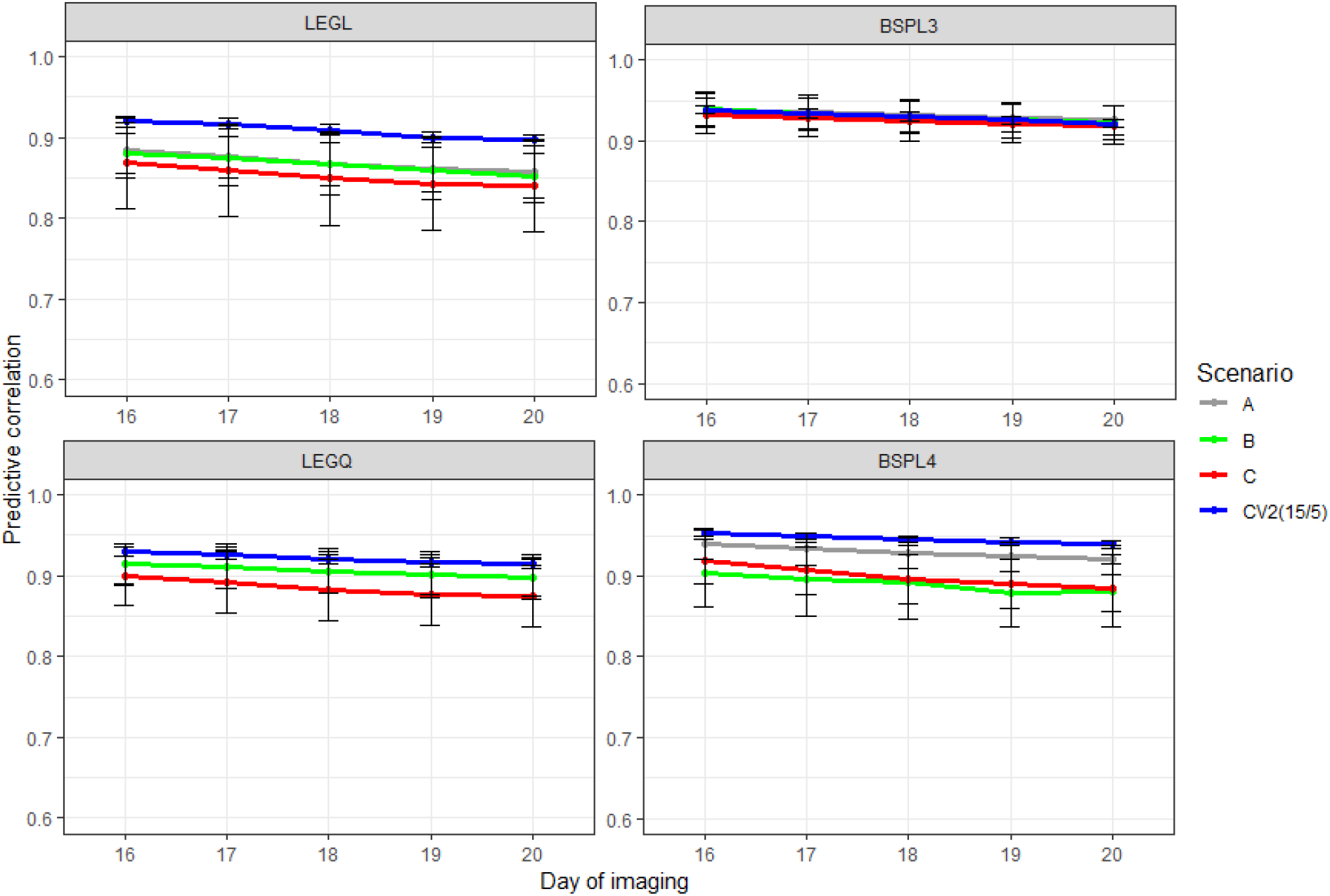
Prediction accuracy of cross-validation scenario 3 in control conditions. A: only observations in the odd days were used; B: only observations in the even days were used; C: keep one and delete two consecutive time points; CV2: use all available previous time points; LEGL: linear Legendre polynomials; LEGQ: quadratic Legendre polynomials; BSPL3: linear B-splines with three knots; BSPL4: linear B-spline with four knots.

**Figure 7:**
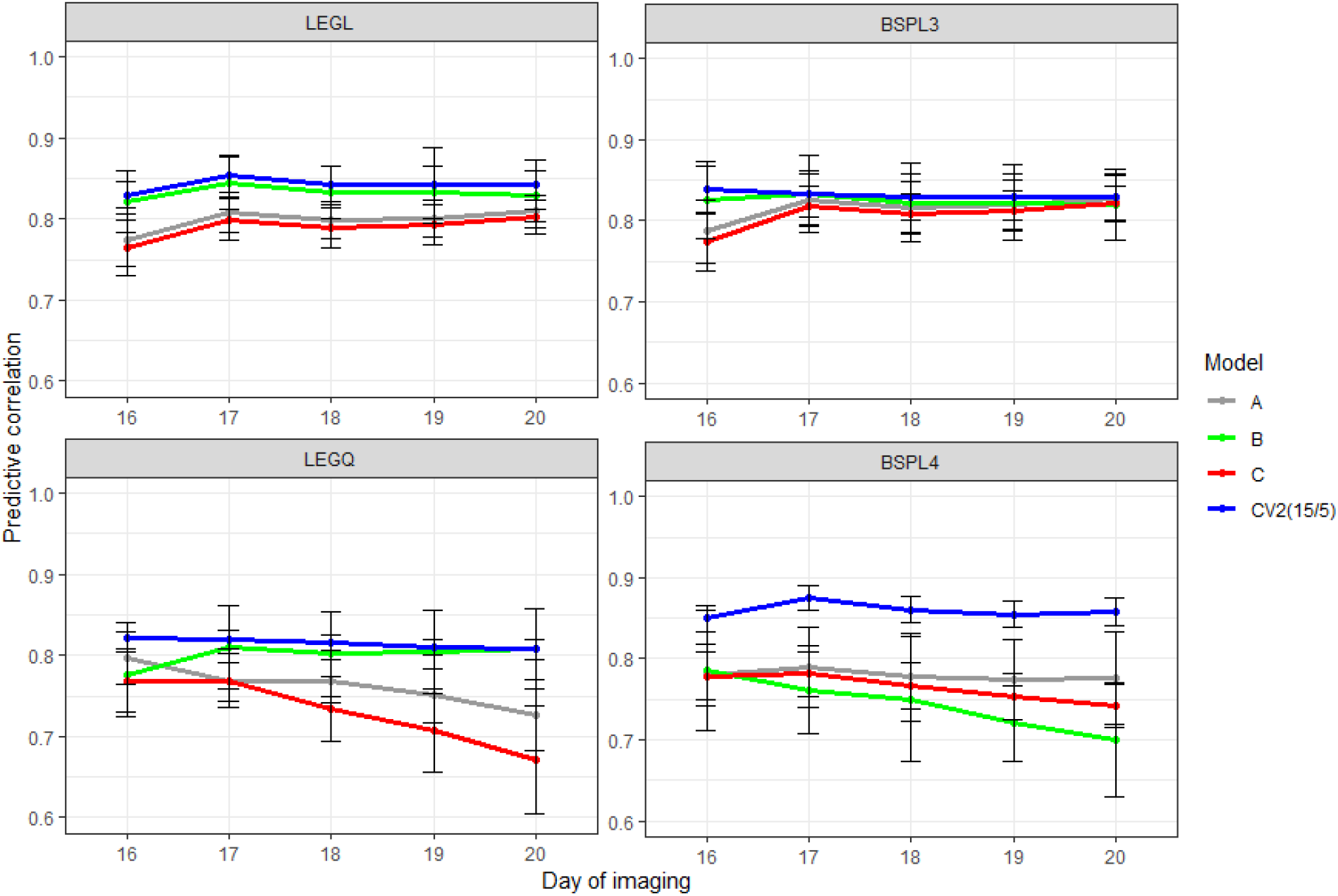
Prediction accuracy of cross-validation scenario 3 in water-limited conditions. A: only observations in the odd days were used; B: only observations in the even days were used; C: keep one and delete two consecutive time points; CV2: use all available previous time points; LEGL: linear Legendre polynomials; LEGQ: quadratic Legendre polynomials; BSPL3: linear B-splines with three knots; BSPL4: linear B-spline with four knots.

## Discussion

Image-based automated HTP technologies offer great potential for characterizing multi-faceted phenotypes at high temporal resolution. The use of HTP platforms plays a pivotal role in accelerating breeding efforts by providing the temporal resolution needed for capturing adaptive responses to environmental challenges, but the development of statistical methodologies to analyze image-based function-valued phenotypes has not kept pace with our ability to generate HTP data. Because phenomics and genomics landscapes for plants are constantly advancing, parallel efforts are required to develop tools for integrating diverse genomic and phenomic datasets characterized by high temporal resolution in genetic analysis. Rice is one of the most drought sensitive cereal crops, resulting in substantial yield losses. With predictions for greater climatic shifts in the future and increasing competition for fresh water resources, research that leverages the full potential of genomics and phenomics is needed to elucidate the genetic and physiological basis of drought tolerance. However, there is currently a lack of information regarding the modeling of temporal HTP data.

RRM identifies the effects of heterogeneous SNPs that transiently influence key traits and translates this to prediction of phenotypes. The main idea behind RRM is to describe subject-specific curves through basis functions (Meyer and Kirkpatrick, 2005). Although RRM has been successfully applied to pedigree-based animal breeding (Schaeffer and Jamrozik, 2008), its utility is largely limited to evaluating goodness-of-fit for candidate models rather than CV-based prediction, and its integration into HTP data has not been reported. In this study, we coupled HTP data with high-density genomic infromation to carry out longitudinal prediction by capturing time-specific genetic signals. A diverse panel of rice accessions subjected to drought stress was used to illustrate the utility of the RRM for evaluating Legendre polynomials and B-splines of time at recording.

### Longitudinal prediction

We found that it was possible to model longitudinal PSA responses under water-limited conditions, albeit with decreased prediction accuracy compared with that of the control. We also placed particular emphasis on comparing two basis functions, i.e., Legendre polynomials and B-splines. To the best of our knowledge, the current study is the first to use a B-spline function to evaluate longitudinal prediction accuracy in the RRM applied to HTP data. Linear B-spline functions with *s* = 3 (two segments) or *s* = 4 knots (three segments) were used. B-splines have been reported to have better numerical properties (e.g., lower computational requirement and faster convergence) than Legendre polynomials because each coefficient of a function affects only a part of the trajectory and can be used to estimate genetic parameters more smoothly while still adequately capturing the features of dynamic data (Iwaisaki et al., 2005; Baldi et al., 2010).

We observed differences in prediction accuracy across models during early growth stages; however, differences were incremental when predicting later growth stages in the CV1 scenario, in which the training and testing sets were partitioned based on individuals. Overall, linear Legendre polynomials performed the best and was clearly an advancement over the MTM. Prediction performance in CV2, in which the training and testing sets were partitioned according to growth stages rather individuals, showed that it was possible to predict future phenotypes from information available from earlier trajectories. Here, linear and quadratic Legendre polynomials produced the highest and most stable prediction accuracy under control conditions, whereas linear B-splines with three knots performed better in the water-limited environment. The final scenario (CV3) demonstrated that we could decrease the phenotyping frequency by only phenotyping every other day to reduce the phenotyping cost while minimizing the loss of prediction accuracy. In this case, linear B-spline with three knots performed relatively well.

B-spline functions require two parameters (the position of the knots and the number of knots) to be tuned. The position of knots can be chosen based on a trajectory pattern such that more knots are placed for rapidly changing time points, whereas less knots are placed for time points with slow changes (Misztal, 2006). Thus, the position of knots should be carefully chosen if the number of phenotyped individuals varies substantially at each growth stage. We chose equidistant knots in the current study because all accessions were phenotyped on the same days during the trajectory. The number of knots determines the number of segments fitted. When more knots are specified, the model becomes more complex. Although we used *s* = 3 and *s* = 4 based on previous literature and a visual inspection of the observed phenotypic trajectory, further investigations are warranted to explore the impact of the number of knots on prediction accuracy. The performance of quadratic B-spline functions was not evaluated in the current study because we encountered convergence issues, possibly due to the small sample size. In general, we found that the advantages of B-splines in inferential tasks compared with Legendre polynomials were not shown clearly in terms of prediction. This is likely because PSA trajectories were relatively simple exponential or monotonically increasing trajectories without obvious inflection points, indicating that the potential of B-splines was not able to be fully exploited in the current study.

### Choice of parameters

We also found that ranking the models according to AIC and BIC revealed only mild agreement with prediction performance evaluated by CV, suggesting that the RRM that gives the best goodness-of-fit is not guaranteed to deliver the best prediction and vice versa. The choice for the order of fit or the number of knots is arguably the most challenging modeling aspect in the RRM. In the majority of literature describing the RRM, this parameter is mainly chosen based on AIC, BIC, or the eigendecomposition of the covariance matrix. The major issue regarding this approach is that there is a tendency to simply pick a model with the highest order of fit or the largest number of knots. However, this study, suggests finding the best parameter in terms of prediction accuracy obtained from CV.

### Future perspective

We anticipate that the current work will guide us to conduct genomic selection of economically important traits on the longitudinal scale for the purpose of breeding crops that are adaptable to new environments or to less favorable challenging climatic conditions. Moreover, identifying genomic components over trajectories will provide information regarding the optimum time points to maximize cost-effective selection or to design a genome-assisted breeding program aiming to change the shape of the longitudinal response curve (Schaeffer, 2004). Using our approach, we could evaluate all changes in plant biomass accumulation during the course of the experiment, in contrast to single time point analyses. Thus, we expect that RRM analysis will become the norm for modeling trajectories of function-valued HTP data because such approaches could be considered an extension of the widely used genomic best linear unbiased prediction model for time series data. Lastly, the utility of the RRM does not preclude its use in other applications. For example, the RRM offers a new avenue for performing longitudinal GWAS (e.g., Howard et al., 2015; Campbell et al., 2019) and genotype-by-environment interactions using the reaction norm (Arnold et al., 2019). In summary, an RRM using Legendre polynomial or spline functions could be an effective option for modeling trait trajectories of HTP data and may have potential applications in characterizing phenotypic plasticity in plants.

## Acknowledgments

This work was supported by the National Science Foundation under Grant Number 1736192 to HW and GM, and Virginia Polytechnic Institute and State University startup funds to GM.

## Author contribution statement

MTC and HW designed and conducted the experiments. MM analyzed the data. MM and GM conceived the idea and wrote the manuscript. MTC and HW discussed results and revised the manuscript. GM supervised and directed the study. All authors read and approved the manuscript.

