## Supplementary material for "Predicting longitudinal traits derived from high-throughput phenomics in contrasting environments using genomic Legendre polynomials and B-splines": Figure S1

### Supplementary Figures

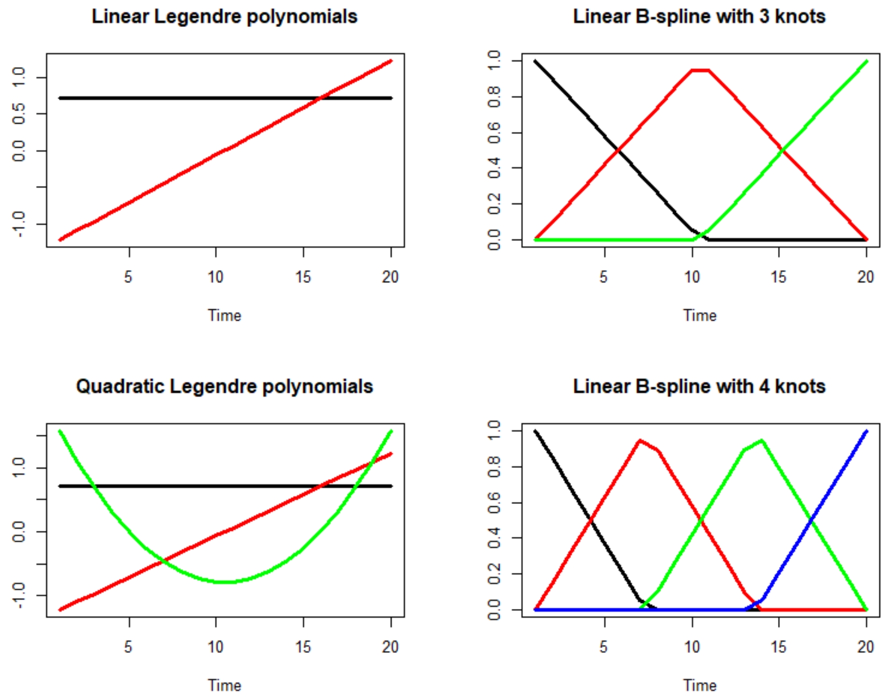


Figure S1: Linear and quadratic forms of Legendre polynomials (left) and three and four knots B-spline curves (right) over 20 days of imaging. Legendre: intercept, linear, and quadratic coefficients are represented in black, red, and green, respectively. B-spline: knot 1, knot 2, knot 3, and knot 4 are represented in black, red, green, and blue respectively.
